# Induction of ectopic external gills and tetrapodomorph-like skeletal elements through homeotic transformations in the salamander branchial region

**DOI:** 10.1101/2024.07.02.601774

**Authors:** Jan Vintr, Vladimír Soukup

## Abstract

Homeotic transformations are morphological changes associated with alterations of identities of segments in serially repeated systems and these changes may be a source of modifications in body plan evolution. Retinoic acid (RA) signaling has previously been shown to induce homeotic transformations in the vertebral column, although its role in other vertebrate segmented systems remains unexplored.

Here, we use pharmacological inhibition of RA receptors to study homeotic transformations in the larval salamander branchial region. This region normally contains three pairs of external gills supported by the underlying skeleton, however upon treatments we observe induction of ectopic outgrowths in the posterior portion of the apparatus. We characterize these outgrowths as ectopically induced fourth external gills on the position of the otherwise gill-free segment. This induction is further associated with transformations and appearance of cartilaginous elements that phenocopy similar elements in fossil stem amphibians and tetrapodomorphs.

These experimentally instigated morphological changes qualify as homeotic transformations in the branchial region and present re-emergence of features that were lost prior to the origin of modern tetrapods. More broadly, our results point to RA signaling as a potent driver regulating the number and composition of pharyngeal segments and thus controlling the evolution of the vertebrate pharyngeal apparatus.

## Introduction

According to the original definition by Bateson (1894), “homeosis” or the nowadays more broadly used term “homeotic transformation” is a phenomenon, whereby “something has been changed into the likeness of something else”. This means that a given structure was transformed into the homologous structure already present on another body segment. Well-known examples of homeosis include transformations of appendages in *Antennapedia* and *Ultrabithorax* fly mutants, transformations in the vertebral column, or identity changes in the floral organs in flowering plants. Vertebrates are special among other organisms in that their body is built based on three independent segmented systems (Graham et al. 2014). Besides segmented somites that give rise to the serially arranged vertebral column and trunk musculature, the vertebrate body plan involves segmentation of a hindbrain into rhombomeres and a pharyngeal region into an apparatus for food intake and gas exchange. The embryonic pharyngeal apparatus is a defining feature of the phylotypic stage of all vertebrates (Irie & Kuratani 2011), including humans. It is composed of arches separated by clefts that undergo evolutionary transformations specific to each vertebrate class. For example, in extant larval salamanders such as the axolotl, this segmented apparatus consists of the mandibular arch forming the jaw, the hyoid arch forming the opercular fold, and the four branchial arches, first three of them bearing the feathery external gills. These structures are further supported by skeletal elements, and so each segment exhibits specific internal and external anatomy (Fig. 1).

**Figure 1.**
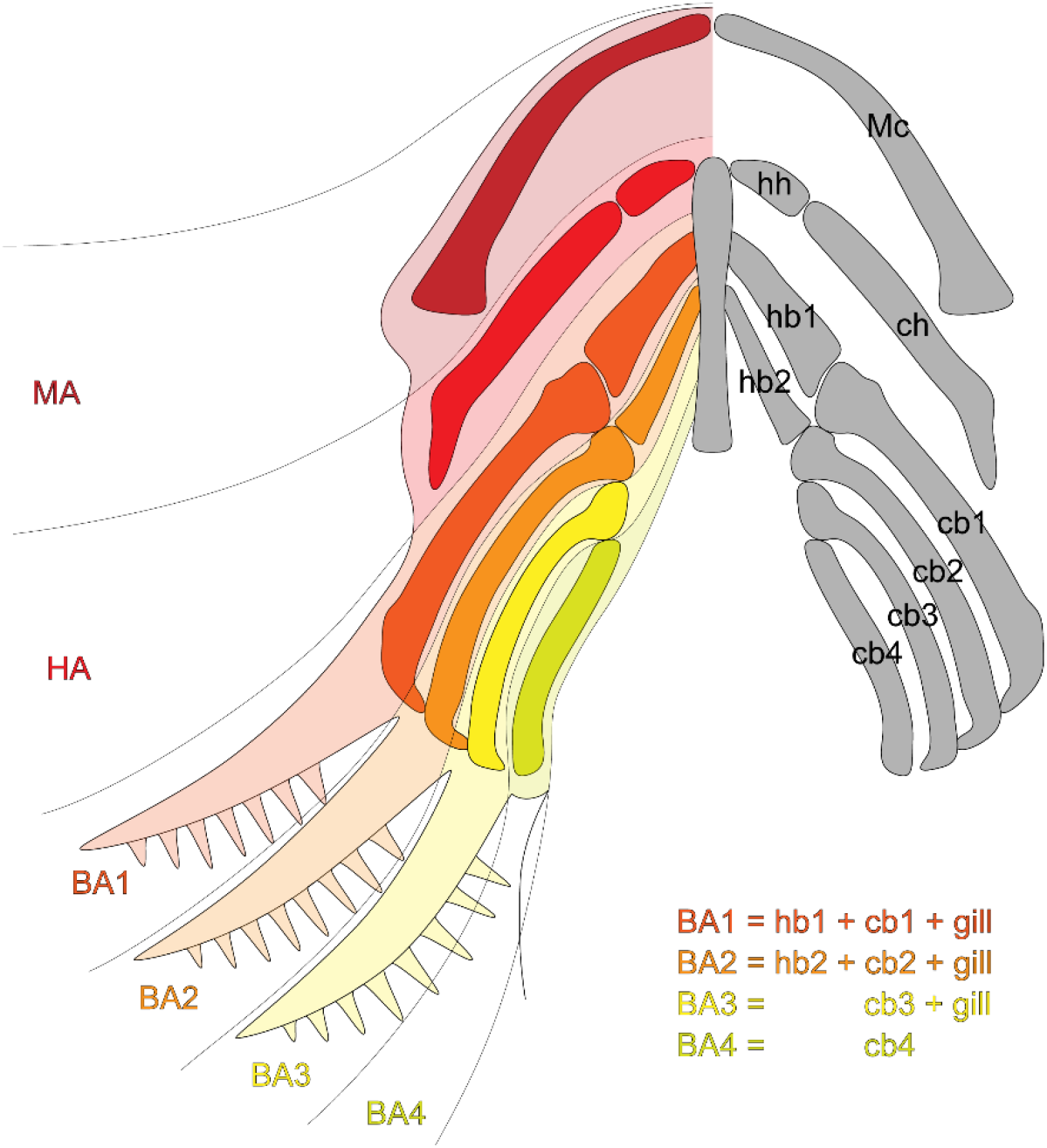
The axolotl pharyngeal apparatus is composed of segments with distinct composition and identities. Major differences are found in the branchial portion of the apparatus. Branchial arches 1 and 2 each contain hypobranchial and ceratobranchial elements and bear an external gill. Branchial arch 3 is different from 1 and 2 in that it lacks a hypobranchial. Branchial arch 4 contains only a ceratobranchial and lacks a hypobranchial as well as an external gill. Abbreviations: BA1-4, branchial arches 1-4; cb1-4, ceratobranchial 1-4; ch, ceratohyal; MA, mandibular arch; Mc, Meckel’s cartilage; HA, hyoid arch; hb1-2, hypobranchial 1-2; hh, hypohyal.

Retinoic acid (RA) signaling has previously been identified as a modulator of identities of segmentally organized structures. In the now classical paper, Kessel & Gruss (1991) showed that exposure of mouse embryos to RA results in homeotic transformations of body segments, affecting the vertebrae and the associated axial skeletal elements. Likewise, previous research in zebrafish has shown that correct development and patterning of the pharyngeal apparatus requires an appropriate amount of RA. For example, loss of *Raldh2*, which encodes the RA production enzyme, leads to skeletal defects and loss of skeletal elements (Begemann et al. 2001). Conversely, blocking endogenous RA degradation induces development of ectopic teeth in the anterior branchial and oral positions, i.e. regions where teeth were once present but became lost during the evolution of cypriniforms including the zebrafish (Seritrakul et al. 2012, Jackman et al. 2024). Tinkering with the amount of RA may thus be a source of variation. However, RA-instigated homeotic transformations have so far not been reported in the branchial region despite an indisputable role of this pathway in its patterning (Mark et al. 2004). We therefore asked if Bateson’s homeotic transformations can be induced in the Mexican axolotl, whose larvae possess distinct and conspicuous identities of individual pharyngeal segments (Fig. 1).

## Results and Discussion

To test this hypothesis, we analyzed expression patterns of *Raldh2* and *Cyp26b1*, whose gene products encode the main retinoic acid producing and inactivating enzymes, respectively. At the time of external gill bulging, *Raldh2* transcripts are predominantly found in the lateral plate mesoderm posteriorly and slightly ventrally to the branchial region (Fig. 2A-C). At stages of external gill outgrowth, strong *Raldh2* expression is still present in the lateral plate mesoderm, but *Raldh2* transcripts appear in ventral gill bases and the outgrowing tips (Fig. 2D-F). Conversely, *Cyp26b1* expression spans the area just dorsally to the outgrowing external gills and the expression is present also on the dorso-anterior portions of individual gills (Fig. 2G-I), suggesting that low concentration of RA is needed for external gills to develop. This pattern is conserved also during the gill outgrowth phase (Fig. 2J-L). Based on this analysis, we assumed that RA signaling may play a role in patterning the antero-posterior axis of the developing branchial region, being produced posteriorly and degraded anteriorly, and potentially having a role in segment identity decisions.

**Figure 2.**
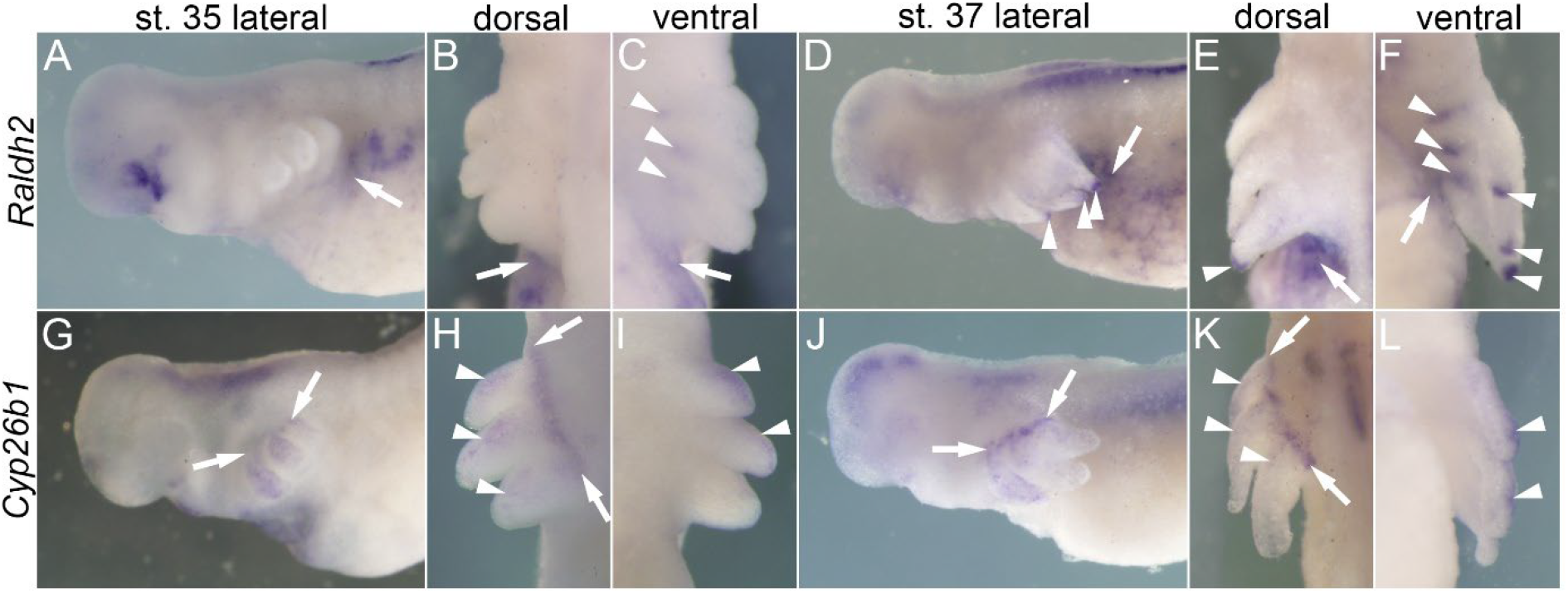
Expression of genes encoding main enzymes responsible for RA production (*Raldh2*) and degradation (*Cyp26b1*). Note the mutually exclusive expression patterns with presence of *Raldh2* transcripts in posteroventral regions behind the gill buds (A, D, arrows) and *Cyp26b1* in anterodorsal regions of gill buds (G, J, arrows). This exclusivity is present also directly on gill buds with *Raldh2* expressed in their ventral portions and the tips (C-F, arrowheads), and *Cyp26b1* in anterior portions of the buds (H, I, K, L, arrowheads).

To test if RA signaling may instigate homeotic transformations in the branchial region, we specifically blocked RAR receptors (but not RXR receptors) using pharmacological agents BMS493, AGN193109, and AGN194310 (Stafford and Prince 2002, Walkley et al. 2022, Hauswirth et al. 2022). Interestingly, upon application of sublethal doses of either one of these pharmacological agents at the time of gill bulging, the treatments resulted in induction of ectopic outgrowths posterior to the three external gills in roughly 70-80% of the cases (Fig. 3A-E). Because of the position of these ectopic outgrowths, we suspected their identity as newly induced external gills. In salamanders, external gills are branched structures composed of the central stalk and minute filaments positioned in two parallel rows along the posterior extent of the stalk (Saito et al. 2019, Ichikawa & Toyoizumi 2020). The gills are composed of stiff connective tissue, and a pair of muscles running and innervated at the level of the filaments. The blood runs in a specific manner from ventral afferent capillaries through the filaments to dorsal efferent capillaries and the gill surface is covered by multi-ciliated cells that increase the flow of the surrounding water to facilitate gas exchange during respiration.

**Figure 3.**
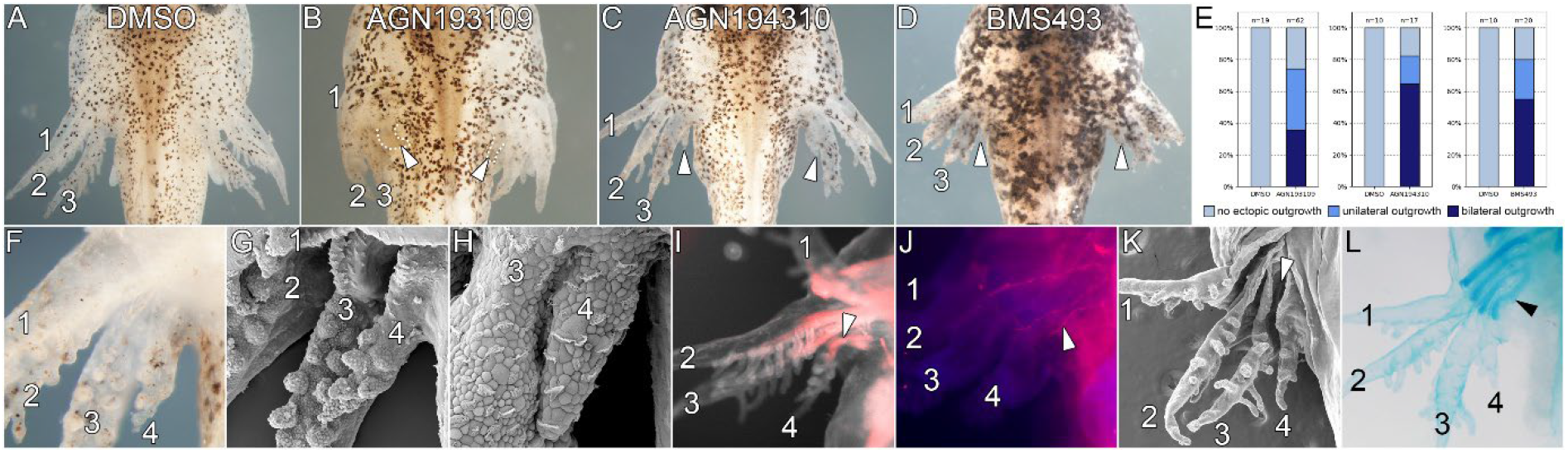
Inhibition of RA receptors leads to induction of ectopic external gills on fourth branchial arches. Ectopic outgrowths are induced posterior to the external gill series upon application of RAR inhibitors AGN193109, AGN194310, and BMS493 (A-E, arrowheads). These outgrowths qualify as ectopic external gills based on the presence of filaments (F-G), multi-ciliated cells (H), musculature (I, arrowhead), and its innervation (J, arrowhead). These ectopic gills develop from the fourth branchial arch that is separated from the third external gill by a gill slit (K, arrowhead). Presence of the fourth gill coincides with altered morphology of the underlying alcian blue-stained cartilaginous visceral skeleton (L, arrowhead).

We analyzed external and internal morphology of the ectopic outgrowths to find that they exhibit gill-like branched external morphology (Fig. 3F). Scanning electron microscopy confirmed the presence of two parallel rows of filaments much like in normal external gills (Fig. 3G). Multiple multi-ciliated cells can be found on the dorsal surface of the outgrowths (Fig. 3H). Antibody staining further confirms presence of muscles and their innervation running along the filament rows (Figs. 3I, J). To demonstrate the presence of vascularization, we captured a timelapse video showing blood circulation in the capillaries much like in normal gills (Supplementary video). Ventral view further shows that this outgrowth is not a mere offshoot of the third external gill but is separated from it by the third gill slit and thus represents a derivative of the fourth branchial arch (Fig. 3K). This analysis thus identifies the outgrowths as ectopically induced external gills in the position of the fourth branchial arch; qualifying this morphological change as a homeotic transformation of the gill-free arch into the gill-bearing arch.

All extant amphibians possess three pairs of external gills. Concordantly, soft tissue imprints of fossil stem tetrapods (such as temnospondyls and seymouriamorphs) confirm this number to be a long-shared amphibian feature (Witzmann 2004). However, in contrast to this evolutionarily conserved gill number, Schmalhausen (1968) reported an extant early branching hynobiid salamander *Ranodon* to possess a rudimentary fourth external gill; and mentions of four-gilled Mexican axolotls are relatively frequent on salamander web fora. It thus seems that fourth gills may occasionally appear in salamanders, suggesting an aberration from the fixed number of external gills.

Whereas all extant amphibians and extinct stem tetrapodomorphs possess only four branchial arches (Witzmann 2013), lungfishes possess four pairs of external gills on branchial arches 1-4 and a gill-free arch 5 (Bartsch et al. 1994, Sefton et al. 2016). In line with our ectopically induced fourth external gills in the axolotl that phenocopy the lungfish-like condition, we inquired if our pharmacological experiments affected the underlying skeleton, potentially inducing development of extra cartilages. In control animals, the pharyngeal skeleton consists of Meckel’s (mandibular) and hyoid cartilages anteriorly, followed by branchial cartilages posteriorly (Fig. 4A). These contain basibranchials 1 and 2, hypobranchials 1 and 2, and ceratobranchials 1-4 (Fig 4B, C).

**Figure 4.**
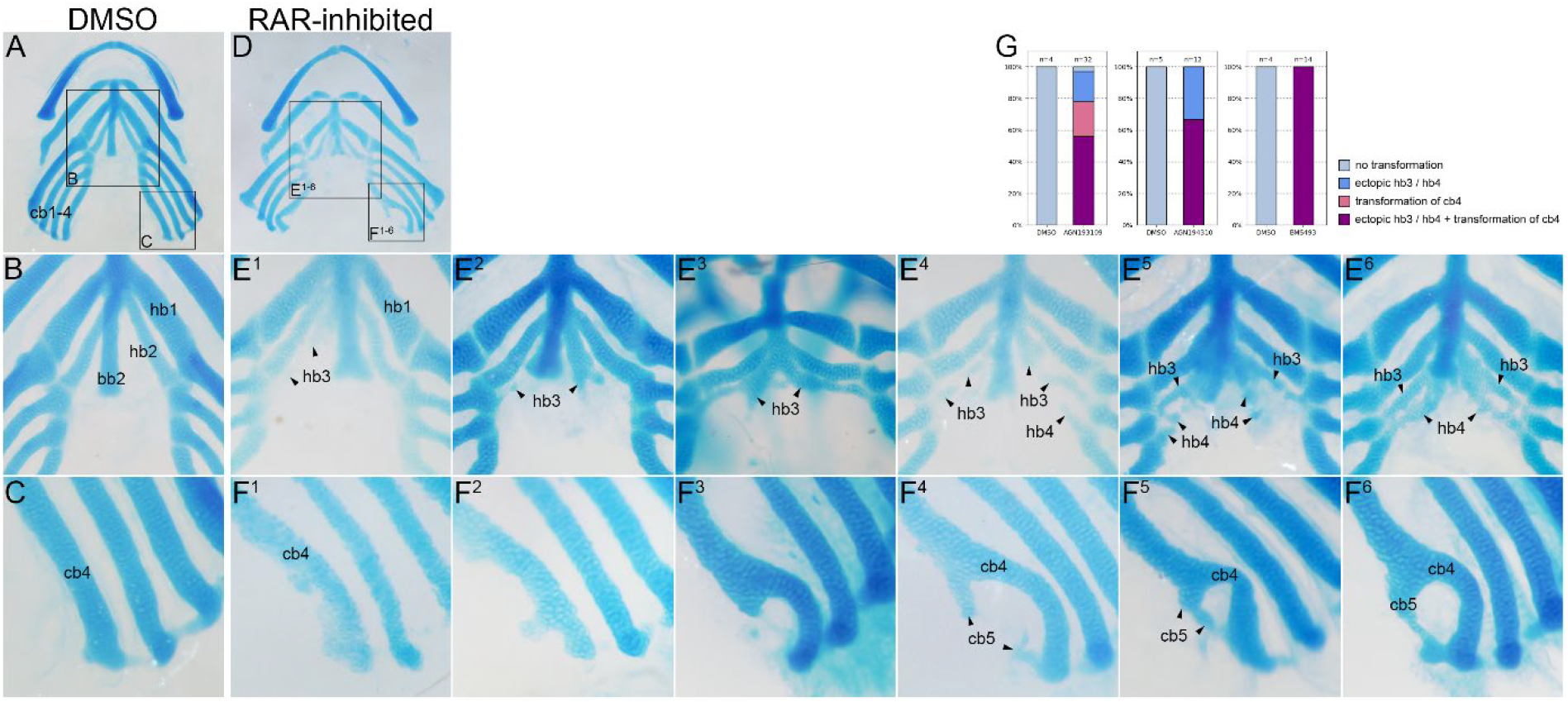
Inhibition of RA receptors results in morphological changes in the visceral skeleton. In DMSO controls, the skeleton contains straight hypobranchial 1-2 and ceratobranchial 1-4 elements (A-C). In RAR-inhibited embryos, the skeleton shows a spectrum of mild-to-severe morphological changes exhibiting induction of extra hypobranchial elements (D, E^1-6^, hb3, hb4), and bifurcation of ceratobranchials 4 into ‘ceratobranchials 5’ (D, F^1-6^, cb5) with high incidence (G). Abbreviations: bb2, basibranchial 2; cb1-4, ceratobranchials 1-4; cb5, ectopic ‘ceratobranchial 5’; hb1-2, hypobranchials 1-2; hb3-4, ectopic ‘hypobranchials 3 and 4’.

In RAR-inhibited animals, the ectopic induction of fourth external gills correlates with altered morphology of the underlying cartilaginous elements (Fig. 3L). These animals exhibit curved morphology of the gill-bearing ceratobranchials 4; this curvature is sometimes subtle, but sometimes forms a complete cartilaginous loop (Fig. 4D, F^1-6^). Although we did not detect any joints, the posterior portion of the loop resembles cartilaginous rod that could be called a newly formed ‘ceratobranchial 5’ (Fig. 4F^5-6^). Moreover, in the median portion of the pharyngeal skeleton, the treated animals exhibit presence of supernumerary elements posterior to and in a sequential order to hypobranchial 2, which we would call ‘hypobranchials 3’ or even ‘hypobranchials 4’ (Fig. 4D, E^1-6^). These morphological changes range from “mild” (Fig. 4E^1-3^, F^1-3^) to “severe” (Fig. 4E^4-6^, F^4-6^), but overall are relatively frequent (Fig. 4G). Thus, in addition to ectopic external gills, the RAR inhibition leads to induction of extra cartilaginous elements posterior to the standard pharyngeal cartilage set.

Interestingly, the RAR-inhibition induced ‘hypobranchial’ elements in the axolotl resemble hypobranchials 3 in members of extant earliest diverging hynobiid and cryptobranchid salamanders (Elwood & Cundal 1994, Deban & Wake 2000, Rose 2003). Moreover, rudimentary hypobranchials 3 may occasionally appear also in salamanders that are normally devoid of them, such as in the fire salamander (*Salamandra salamandra*), the spotted salamander (*Ambystoma maculatum*), or the Eastern newt (*Notophthalmus viridiscens*) (Kallius 1901, Rose 2003, Gao & Shubin 2012), and rudimentary hypobranchials 4 were even noted in the cryptobranchid hellbender (*Cryptobranchus alleganiensis*) (Drüner 1904). Even more interestingly, the RAR-induced ‘hypobranchial’ elements in the axolotl resemble hypobranchials 3 of the Paleozoic stem tetrapods such as *Adelogyrinus, Gerrothorax*, and *Trimerorhachis* (Witzmann 2013), and hypobranchials 3 and 4 of other stem tetrapods and finned tetrapodomorphs *Brachydectes, Mandageria, Eusthenopteron, Glyptolepis, Gogonasus*, and *Tiktaalik* (Jarvik 1980, Long et al. 1997, Downs et al. 2008, Johanson & Ahlberg 1997, Witzmann 2013). Besides the hypobranchials, the experimentally induced ‘ceratobranchials 5’ resemble the basic condition of five branchial arches shared among all bony vertebrates (Osteichthyes) and found in extant lungfishes and the coelacanth (Bartsch 1994, Sefton et al. 2016). Our RAR-inhibition treatments in axolotl thus phenocopy the primitive skeletal composition of tetrapod ancestors found about 140-350 Mya that predates the derived reduced condition of modern salamanders (Gao & Shubin 2012).

Our experimentally induced atavistic appearance of external gills and pharyngeal skeletal elements together with the previously described occasional appearance of these morphological features in various salamander species hints at RA signaling as a potential key driver of the number and composition of the posterior pharyngeal arches. This regulation may even predate vertebrates as RA signaling regulates the posterior extent of the pharynx in the cephalochordate amphioxus by repressing the expression of pharynx-specific genes (Schubert et al. 2001). Tinkering with the amount of RA may thus be a generic source of variability of pharyngeal apparatuses among chordates and our data on transformations of axolotl posterior branchial arches are well in line with this supposition. In addition, experimental induction of fourth external gills together with the underlying branchial skeleton in the axolotl represents, to our knowledge, the first case of homeotic transformation induced in the branchial region and shows the power of the salamander pharyngeal apparatus as a tractable segmented system, where traditional Bateson’s homeoses can eligibly be studied.

## Acknowledgments

The authors thank Ann Huysseune for many discussions on the topic, and Himanshi Singh, Jan Stundl and Marianne E. Bronner for critically reading earlier versions of the manuscript. The authors further thank the Tanaka lab (IMP Vienna) for providing the *Raldh2* clone. This project was supported by the Grant Agency of the Charles University GAUK 355022 (to J.V.), the Czech Science Foundation GACR 23-07212S, and Charles University Research Center Program 204069 (both to V.S.). The authors acknowledge Vinicna Microscopy Core Facility of the Faculty of Science, Charles University, an institution supported by the MEYS CR (LM2023050 Czech-BioImaging), for their support and assistance in this work.

## Author contributions

JV and VS designed research; JV performed research; JV analyzed data; VS wrote first draft of the manuscript; both authors read and approved final version of the manuscript.

## Supplementary Information

### Experimental procedures

#### Animal handling and pharmacological inhibitions

Melanic and albino embryos of the Mexican axolotl were acquired from the colony at the Department of Zoology, Charles University, Prague. Embryos were manually dechorionated and staged according to Bordzilovskaya et al. (1989). Pharmacological compounds BMS493 (Sigma B6688), AGN193109 (Sigma SML2034), and AGN194310 (Sigma SML2665) were dissolved in DMSO to stock solutions of 5 mM, 5 mM, and 4 mM, respectively, and stored in -80°C. Embryos were treated with effective concentrations 5–17.5 µM BMS493, 12.5 µM AGN193109, and 0.8–1.6 µM AGN194310 in 10 ml 1× Steinberg solution. Embryos were treated from stage 35 (gill bulging stage) for about 12 days for subsequent morphological analyses.

#### Gene cloning and whole mount in situ hybridization

Partial sequence of the axolotl *Cyp26b1* was cloned based on the sequence AMEX60DD301046276.2 from axolotl-omics.org using following primers: forward 5’-GGCAACGGTATCCTCCACAA-3’, reverse 5’-TTTCCATTCTTCCCTGCCCC-3’. The PCR product was cloned in pGEM T-Easy to synthesize DIG-labeled probe for in situ hybridization. The axolotl *Raldh2* clone was a gift from the Tanaka lab. Whole mount in situ hybridization was performed as described (Šimková et al. 2023).

#### Whole mount antibody staining

Whole mount antibody staining was performed using similar protocol as described in Smith & Armstrong (1990). Samples stored in methanol were washed in PBS + 0.4% Triton and incubated in primary antibodies in 4% BSA in PBS + 0.4% Triton. The gill muscles were stained with 12/101 antibody (DSHB, AB_531892) (Ericsson & Olsson 2004), and their innervation with anti-myelin basic protein (GeneTex, GTX76114) (Pende et al. 2020). Following incubation at 4°C for 2-10 days, the samples were washed with PBS + 0.4% Triton and incubated in secondary antibodies overnight. Washed samples were photographed using Zeiss SteREO Lumar.V12 fluorescent stereomicroscope.

#### Alcian blue staining

Samples stored in methanol were stained in alcian blue solution (80% acetic acid + 20% ethanol, 0.015 g alcian blue per 100 ml solution) for at least 48 hours. Samples were then bleached in 0.75% H_2_O_2_ and either directly cleared in 0.5% KOH + glycerol (1:1) for at least 7 days or treated with 0.3% trypsin and then cleared. Upon dissection at the level of the jaw joint, the released Meckel’s cartilage and the hyobranchial apparatus were photographed using Olympus SZX12 stereomicroscope.

#### Scanning electron microscopy

Samples fixed in 4% PFA were dehydrated in ethanol, desiccated in Bal-Tec CPD 030 critical point dryer, coated in gold using Bal-Tec SCD 050 sputter coater, and attached onto carbon adhesive tabs. SEM microscopy was performed using JEOL JSM-6380 LV scanning electron microscope.

## References

Bartsch P (1994) Development of the cranium of Neoceratodus forsteri, with a discussion of the suspensorium and the opercular apparatus in Dipnoi. Zoomorph 114, 1–31.

Bateson W (1894) Materials for the study of variation treated with especial regard to discontinuity in the origin of species. Macmillan, London. 598 p.

Begeman G, Schilling TF, Rauch GJ, Geisler R & Ingham PW (2001) The zebrafish neckless mutation reveals a requirement for raldh2 in mesodermal signals that pattern the hindbrain. Development 128, 3081–3094.

Deban SM & Wake DB (2000) Aquatic feeding in salamanders. In: Feeding. Form, Function and Evolution in Tetrapod Vertebrates (Schwenk K, ed.), Academic Press, San Diego, pp. 65–94.

Downs JP, Daeschler EB, Jenkins FAJr & Shubin NH (2008) The cranial endoskeleton of Tiktaalik roseae. Nature 455, 925–929.

Drüner L (1905) Studien zur Anatomie der Zungenbein-, Kiemenbogenund Kehlkopfmuskeln der Urodelen. II Theil. Zool Jahrb 19, 361–690.

Elwood JRL & Cundal D (1994) Morphology and behavior of the feeding apparatus in Cryptobranchus alleganiensis (Amphibia: Caudata). J Morph 220, 47–70.

Gao KQ & Shubin NH (2012) Late Jurassic salamandroid from western Liaoning, China. Proc Nat Acad Sci USA 109, 5767–5772.

Graham A, Butts T, Lumsden A & Kiecker C (2014) What can vertebrates tell us about segmentation? EvoDevo 5, 24.

Hauswirth GM, Garside VC, Wong LSF, Bildsoe H, Manent J, Chang YC, Nafzger CM, Firas J, Chen J, Rossello FJ, Polo JM & McGlinn E (2022) Breaking constraint of mammalian axial formulae. Nat Comm 13, 243.

Ichikawa R & Toyoizumi R (2020) Finely tuned ciliary alignment and coordinated beating generate continuous water flow across the external gills in Pleurodeles waltl larvae. Zoomorph 139, 247–262.

Irie N & Kuratani S (2011) Comparative transcriptome analysis reveals vertebrate phylotypic period during organogenesis. Nat Comm 2, 248

Jackman WR, Portillo LSM, Cox CK, Ambrosio A & Gibert Y (2024) Blocking endogenous retinoic acid degradation induces oral tooth formation in zebrafish. Proc Nat Acad Sci USA 121, e2321162121.

Jarvik E (1980) Basic structure and evolution of vertebrates, Volume 1. London: Academic Press.

Johanson Z & Ahlberg PE (1997) A new tristichopterid (Osteolepiformes: Sarcopterygii) from the Mandagery Sandstone (Late Devonian, Famennian) near Canowindra, NSW, Australia. Trans R Soc Edinburgh Earth Sci 88, 39–68.

Kallius E (1901) Beiträge zur Entwickelung der Zunge. 1. Teil. Amphibien und Reptilien. Anat Hefte 16, 531–760.

Kessel M & Gruss P (1991) Homeotic transformations of murine vertebrae and concomitant alteration of Hox codes induced by retinoic acid. Cell 67, 89–104.

Long JA, Barwick RE & Campbell KSW (1997) Osteology and functional morphology of the osteolepiform fish Gogonasus andrewsae Long, 1985, from the Upper Devonian Gogo Formation, Western Australia. Rec West Austral Mus Suppl 53, 1–89.

Mark M, Ghyselinck NB & Chambon P (2004) Retinoic acid signalling in the development of branchial arches. Curr Opin Gen Dev 14, 591–598.

Rose CS (2003) The developmental morphology of salamander skulls. In: Amphibian Biology Vol. 5 Osteology (eds. H Heatwole, M Davies), Australia: Surrey Beatty and Sons Pty. Ltd., pp. 1686–1783.

Saito N, Nishimura K, Makanae A & Satoh A (2019) Fgf- and Bmp-signaling regulate gill regeneration in Ambystoma mexicanum. Dev Biol 452, 104–113.

Schmalhausen II (1968) The origin of terrestrial vertebrates. Academic Press, 314 p.

Schubert M, Yu JK, Holland ND, Escriva H, Laudet V & Holland LZ (2001) Retinoic acid acts via Hox1 to establish the posterior limit of the pharynx in the cephalochordate amphioxus. Development 132, 61–73.

Sefton EM, Bhullar BAS, Mohaddes Z & Hanken J (2012) Evolution of the head-trunk interface in tetrapod vertebrates. eLife 5, e09972.

Seritrakul P, Samarut E, Lama TTS, Gibert Y, Laudet V & Jackman WR (2012) Retinoic acid expands the evolutionarily reduced dentition of zebrafish. FASEB J 26, 5014–5024.

Stafford D & Prince VE (2002) Retinoic acid signaling is required for a critical early step in zebrafish pancreatic development. Curr Biol 12, 1215–1220.

Walkley CR, Yuan YD, Chandraratna RAS & McArthur GA (2002) Retinoic acid receptor antagonism in vivo expands the numbers of precursor cells during granulopoiesis. Leukemia 16, 1763–1772.

Witzman F (2004) The external gills of Paleozoic amphibians. N Jb Geol Paläont Abh 232, 375–401.

Witzman F (2013) Phylogenetic patterns of character evolution in the hyobranchial apparatus of early tetrapods. Trans R Soc Edinburgh Earth Env Sci 104, 145–167.

## Supplemental information references

Bordzilovskaya NP, Detlaff TA, Duhon ST & Malacinski GM (1989) Developmental-stage series of axolotl embryos. In: Developmental Biology of the Axolotl (ed. Malacinski GM). Oxford University Press, Oxford, 201–219.

Ericsson R & Olsson L (2004) Patterns of spatial and temporal visceral arch muscle development in the Mexican axolotl (Ambystoma mexicanum). J Morph 261, 131–140.

Pende M, Vadiwala K, Schmidbaur H, Stockinger AW, Murawala P, Saghafi S, Dekens MPS, Becker K, Revilla-i-Domingo R, Papadopoulos SC, Zurl M, Pasierbeck P, Simakov O, Tanaka EM, Raible F & Dodt HU (2020) A versatile depigmentation, clearing, and labeling method for exploring nervous system diversity. Sci Adv 6, eaba0365.

Šimková K, Naraine R, Vintr J, Soukup V & Šindelka R (2023) RNA localization during early development of the axolotl. Front Cell Dev Biol 11, 1260795.

Smith SC & Armstrong JB (1990) Whole-mount immunocytochemistry in axolotl embryos. Ax Newsletter 19, 28–30.

